# ChemGenXplore: An Interactive Tool for Exploring and Analysing Chemical Genomic Data

**DOI:** 10.1101/2025.08.13.670066

**Authors:** Huda Ahmad, Hannah M Doherty, Sam Benedict, James Haycocks, Ge Zhou, Patrick Moynihan, Danesh Moradigaravand, Manuel Banzhaf

**Author notes:** Correspondence should be address to: Manuel Banzhaf, Danesh Moradigaravand, Huda Ahmad.

## Abstract

**Motivation:** Chemical genomics is a useful high-throughput approach to systematically link phenotypes to genotypes. However, the vast datasets generated remain challenging to explore due to the lack of integrated, interactive tools for visualisation and analysis. Existing workflows often require multiple independent software tools, limiting data accessibility and collaboration. Therefore, we created a user-friendly platform that enable efficient exploration and sharing of chemical genomics data.

**Results:** We developed ChemGenXplore, a web-based Shiny application designed to streamline the visualisation and analysis of chemical genomic screens. It offers two primary functionalities: one for exploring pre-implemented datasets and another for analysing user-uploaded datasets. ChemGenXplore enables users to visualise phenotypic profiles, assess gene-gene and condition-condition correlations, perform GO and KEGG enrichment analysis, and generate customisable, interactive heatmaps. By consolidating these features into an interactive and accessible tool, ChemGenXplore facilitates data sharing, enhances reproducibility, and promotes collaboration within the research community.

**Availability and Implementation:** ChemGenXplore is freely accessible as a web application at https://chemgenxplore.kaust.edu.sa/ and is also available for local deployment via GitHub at https://github.com/Hudaahmadd/ChemGenXplore.

## 1. Introduction

Over recent decades, advances in high-throughput technologies have facilitated the generation of large-scale molecular and phenotypic data across a wide range of microbial species. These advancements have led to gene expression studies across thousands of conditions (Brown et al., 2014, Chen et al., 2018, Gerstein et al., 2014, Tsankov et al., 2015), as well as genome-wide phenotypic profiling discovering new biology (Arend et al., 2016, Bobonis et al., 2022, Fajardo et al., 2008, Gomez Maria and Neyfakh Alexander, 2006, Koo et al., 2017, Nichols et al., 2011, Schuldiner et al., 2005, Tamae et al., 2008, Tong et al., 2001, Typas et al., 2010). However, the rapid accumulation and complexity of high-throughput biological data present significant computational challenges, necessitating the development of efficient tools for data integration, processing, and interpretation (Houle et al., 2010, Schadt et al., 2010, Sullivan et al., 2014). As high-throughput approaches continue to expand, addressing these challenges is essential for translating large-scale biological data into knowledge.

Chemical genomics is a high-throughput screening method that facilitates the functional annotation of orphan genes by systematically evaluating the effect of gene disruption on fitness across diverse chemical and environmental perturbations (Cain et al., 2020). This approach generates complex phenotypic profiles that serve as a valuable resource for future investigations into gene function and cellular responses (Alodaini et al., 2024, Cain et al., 2020, Doherty et al., 2023, Hayward et al., 2025, Nichols et al., 2011, Schuldiner et al., 2005). Such studies have contributed to the construction of genotype-phenotype relationships and gene interaction networks (Koo et al., 2017, Nichols et al., 2011, Schuldiner et al., 2005, Tong et al., 2001). These experiments involve thousands of gene knockouts, generated through methods such as transposon insertion sequencing (Tn-seq) or deletion libraries, and each mutant is tested across hundreds of stress conditions (Koo et al., 2017, Kritikos et al., 2017, Nichols et al., 2011, Williams et al., 2025). Such datasets cannot be manually analysed to fully capture the phenotypic profile of an organism. In response to these large-scale datasets, a variety of advanced analysis approaches have emerged (Collins et al., 2006, Dixon et al., 2009, Doherty et al., 2023, French et al., 2016, Nichols et al., 2011, Shiver et al., 2017). These approaches provide systematic methods to analyse and quantify phenotypic changes, generating scored datasets. However, the key challenge lies in extracting meaningful biological insights from these scores. Many current pipelines rely on multiple independent tools, such as Cluster 3.0 (de Hoon et al., 2004) for hierarchical clustering, Java TreeView (Saldanha et al., 2004) for data visualisation, in-house scripts, and manual analysis to understand phenotypic profiles. These tools require extensive preprocessing, manual parameter selection, and frequent transitions between platforms, making data exploration inefficient and time-consuming.

Furthermore, the fragmented nature of existing resources limits the accessibility and reproducibility of chemical genomics data, posing barriers to collaborative research.

To address these challenges and facilitate more accessible exploration of chemical genomics data, we introduce ChemGenXplore. ChemGenXplore is a web-based, interactive platform that enables users to visualise phenotypic data, compute gene-gene and condition-condition correlations, perform Gene Ontology (GO) and Kyoto Encyclopedia of Genes and Genomes (KEGG) enrichment analyses, and generate customisable interactive heatmaps with clustering options for genes and conditions. Additionally, ChemGenXplore serves as a user-friendly tool for sharing chemical genomics resources, enhancing accessibility, reproducibility, and collaboration within the field.

## 2. Materials and methods

### 2.1 Overview

ChemGenXplore is designed to facilitate the exploration and interpretation of high-throughput chemical genomic screens. Figure 1 provides a schematic overview of the chemical genomic screening workflow and the role of ChemGenXplore in the data analysis pipeline. In a typical screen, mutant or microbial strain libraries are arrayed onto agar media containing specific chemical stressors. Following incubation, a phenotype such as colony growth is measured through high-throughput imaging, and image analysis pipelines are used to quantify strain fitness. Once the fitness scores are computed, ChemGenXplore enables users to interactively visualise, filter, and interpret the dataset, supporting downstream analyses such as phenotype profiling, correlation mapping, enrichment analysis, and heatmap clustering. By integrating these features into a single interface, ChemGenXplore streamlines the interpretation of complex chemical-genomic data and enhances biological insight into gene- and condition-specific phenotypes.

**Figure 1.**
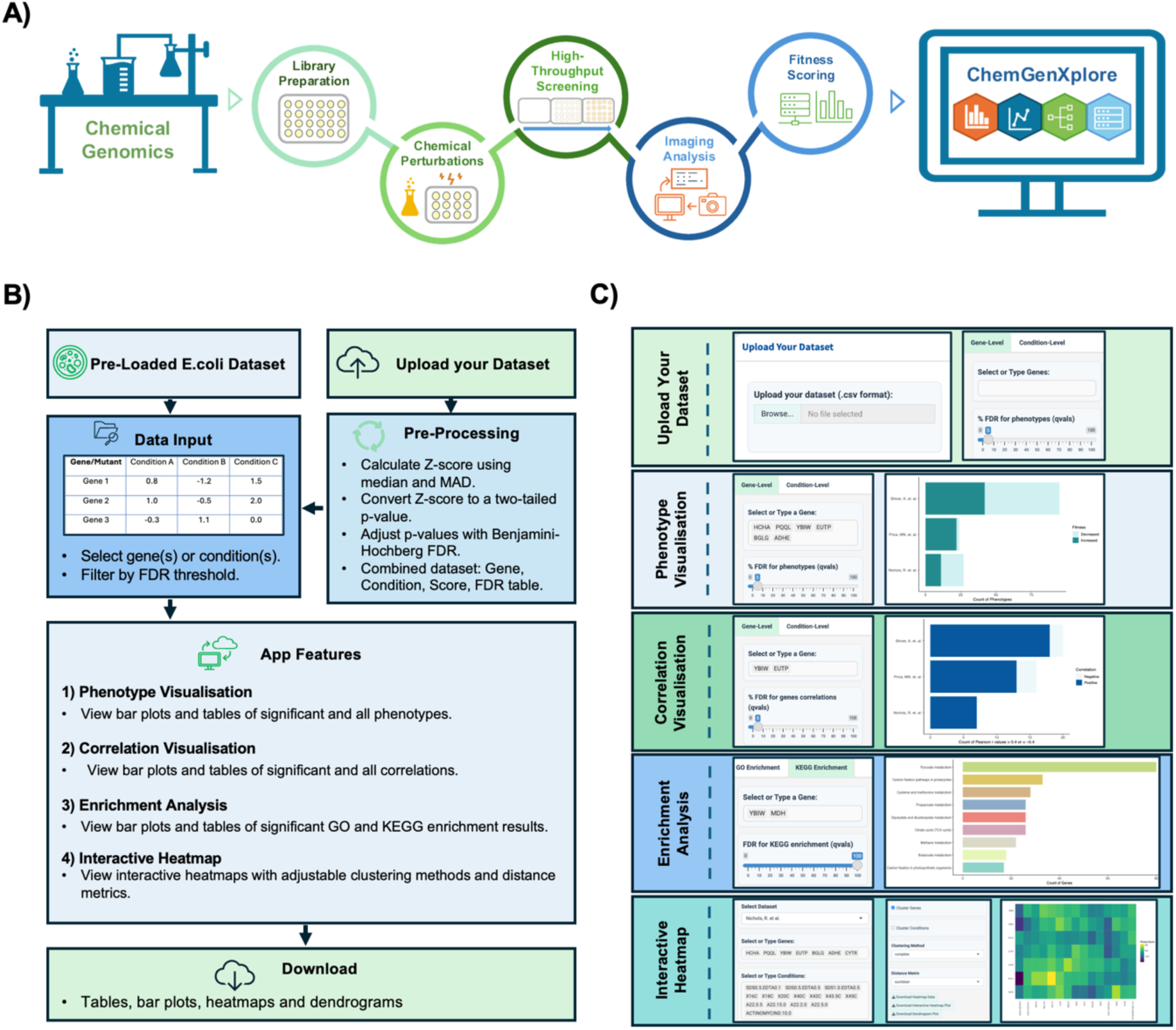
General workflow of chemical genomic screening. **A)** Mutant or microbial strain libraries are arrayed onto plates with chemical perturbations, followed by high-throughput screening, image analysis, and fitness scoring. ChemGenXplore is then used for data visualisation and exploration of the scored dataset**. B)** In ChemGenXplore Users select gene(s) and/or condition(s) of interest to explore phenotypic scores via bar plots and data tables. Other features include gene-gene or/and condition-condition correlation analysis, GO and KEGG enrichment, and interactive heatmaps. Users can also upload their own datasets and export all visualisations and results. **C)** ChemGenXplore Outputs examples from ChemGenXplore: interface panels and plots from each feature.

### 2.2 Data Input

ChemGenXplore allows the upload of multiple datasets, each with a file size limit of up to 10 GB, to accommodate the large-scale nature of chemical genomic screens. The current implementation includes three pre-integrated, publicly available *Escherichia coli* datasets from chemical genomic studies (Fig. 2) (Nichols et al., 2011, Price et al., 2018, Shiver et al., 2017). These datasets provide fitness scores under a wide range of conditions, forming the foundation for the analyses performed in ChemGenXplore. To ensure compatibility, input datasets must adhere to specific formatting requirements. Uploaded datasets should be in CSV format. Each dataset is expected to be in tabular form, where rows represent strains or genes identifiers, and columns correspond to experimental conditions, along with associated phenotypic measurements such as fitness scores.

**Figure 2.**
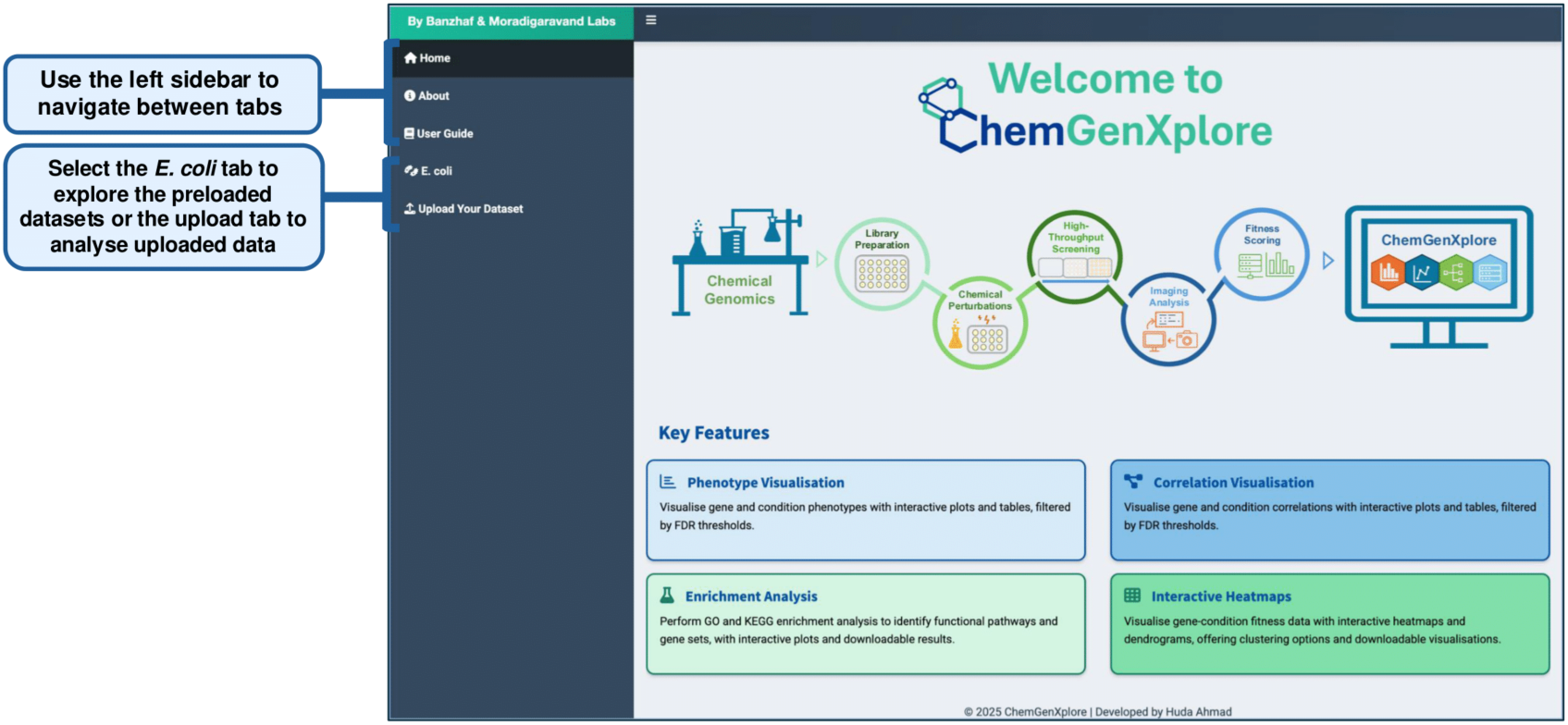
Data Input. ChemGenXplore supports both pre-implemented *E. coli* datasets and user-uploaded datasets.

### 2.3 ChemGenXplore Web Application

ChemGenXplore was developed as an interactive Shiny application using R (https://www.R-project.org/). The web application is freely accessible at: https://chemgenxplore.kaust.edu.sa/. The source code is publicly available at: https://github.com/Hudaahmadd/ChemGenXplore.

## 3. Results

ChemGenXplore is designed to facilitate the interactive exploration of chemical genomic datasets, providing a flexible platform for analysing gene- and condition-specific phenotypes. Figure 3 presents a summary of the functionalities of ChemGenXplore. In the following sections, we outline these functionalities in detail.

**Figure 3.**
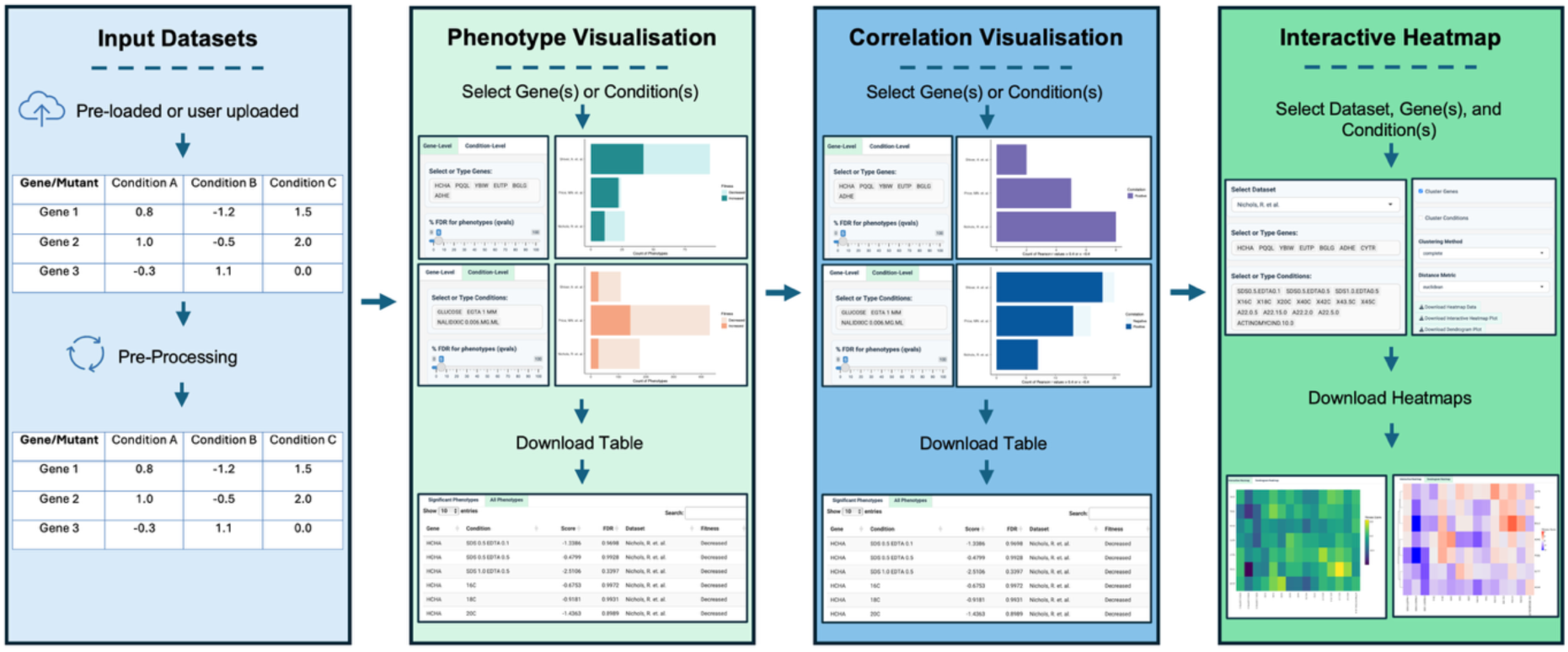
Basic functionality of ChemGenXplore. ChemGenXplore supports the visualisation and analysis of chemical genomic datasets through four key features. Users can select pre-loaded datasets or upload their own in CSV format. Uploaded dataset is automatically pre-processed. The Phenotype Visualisation allows users to explore fitness scores by selecting gene(s) or condition(s) of interest. The Correlation Visualisation enables the identification of gene-gene and condition-condition correlations. The Interactive Heatmap facilitates the clustering of selected genes and conditions based on phenotypic profiles. Each feature supports interactive exploration and the option to download plots and tables for further analysis.

### 3.1 Dataset Upload

ChemGenXplore provides users with the ability to upload their own datasets, where rows represent unique identifiers (e.g., genes, samples, strains) and columns contain experimental measurements (e.g., fitness scores, gene expression levels, growth rates) (Fig. 4). Once a dataset is uploaded, users can perform all the analyses available in the app, including phenotype visualisation, correlation analysis, and heatmap generation. This feature makes it suitable for a wide range of chemical genomic datasets.

**Figure 4.**
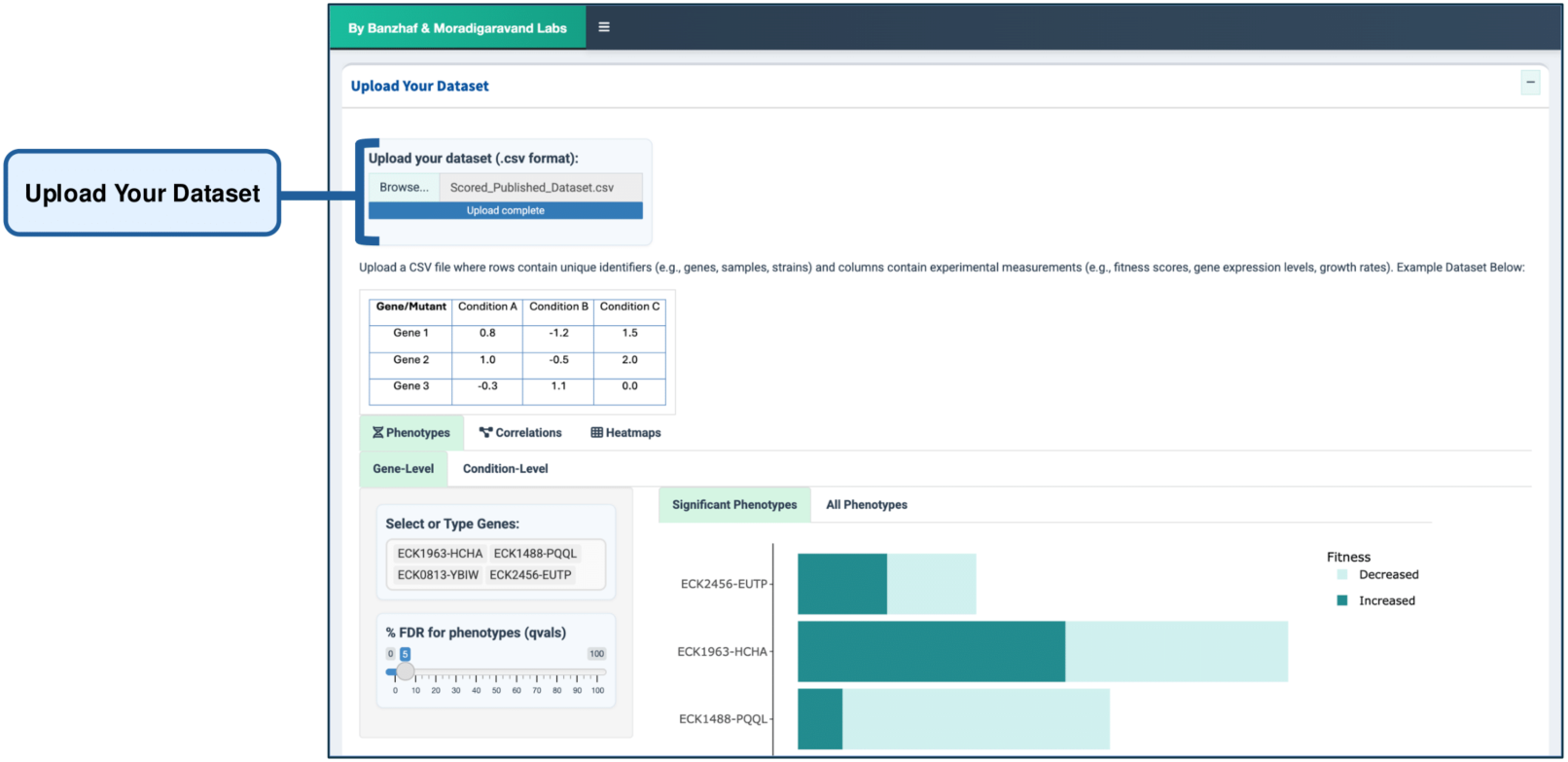
Dataset Upload. Users can upload their datasets in CSV format, where rows represent genes/mutants and columns correspond to conditions. Once uploaded, the dataset is processed, allowing users to perform all available analyses, including phenotype visualisation with FDR analysis, correlation analysis, and heatmap generation, following the same options as in the pre-implemented datasets.

### 3.2 Phenotype Visualisation

The Phenotype Visualisation feature enables users to explore gene- and condition-specific phenotypes through interactive bar plots and data tables. As shown in Figure 5, users can select genes or conditions of interest and view corresponding phenotypic scores, with the option to filter results based on the False Discovery Rate (FDR) threshold. The FDR ensures statistical robustness by controlling false positives, reducing the likelihood of misleading associations that could mask true genotype-phenotype relationships. This feature is particularly useful for highlighting significant phenotypic changes and identifying potential gene-phenotype relationships.

**Figure 5.**
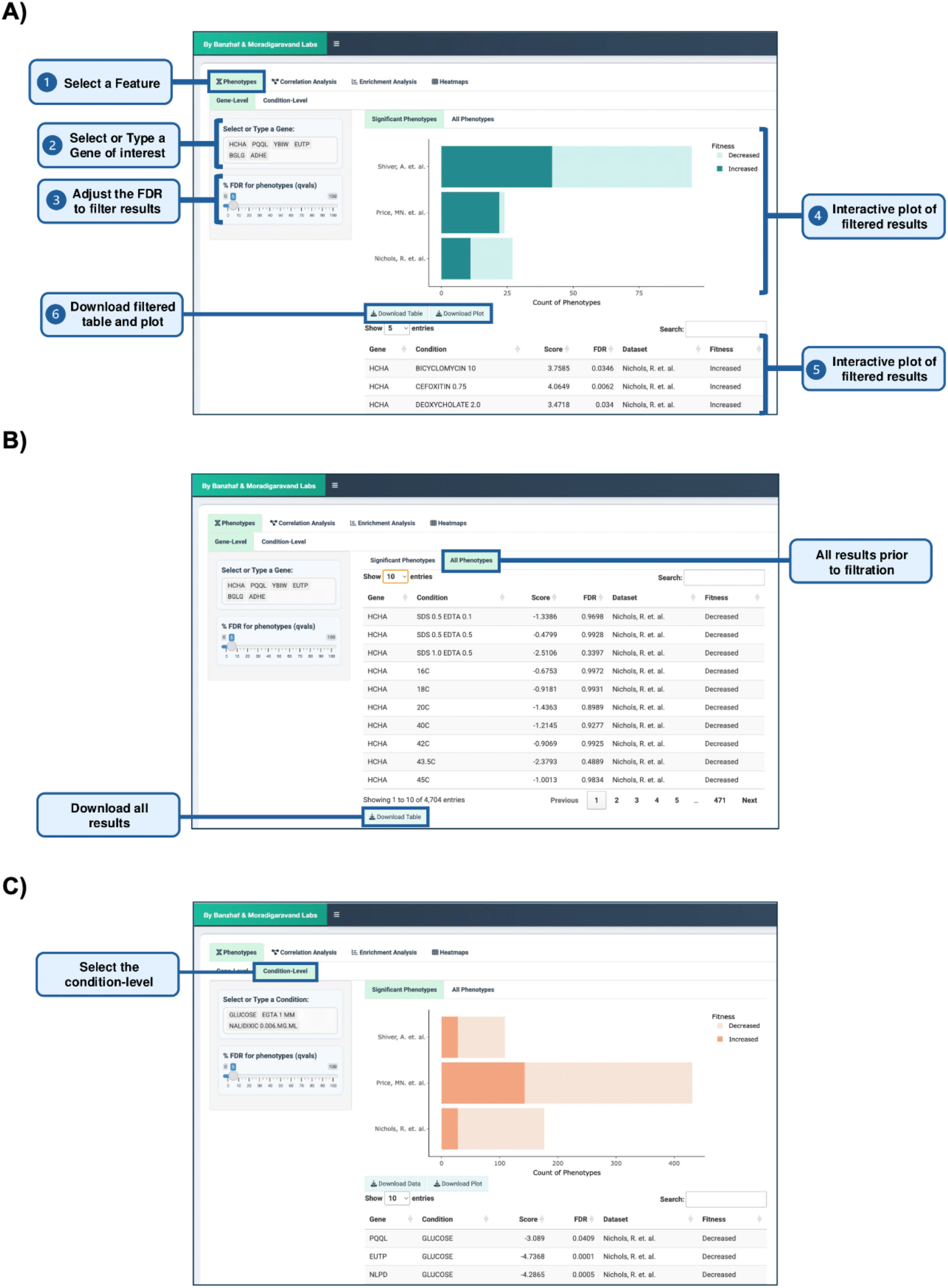
Phenotype Visualisation. **(A)** Users can select or type a single gene or multiple genes of interest to display phenotypes through interactive bar plots and data tables, with an FDR threshold to filter significant results. Both plots and tables are downloadable. **(B)** The "All Phenotypes" tab allows users to view all phenotypes related to the selected gene(s) without filtering. **(C)** Users can explore condition-specific phenotypes with the same display and download options.

### 3.3 Correlation Analysis

Correlation analysis is a fundamental approach in chemical genomics for identifying functionally related genes and condition-specific phenotypic patterns. High correlation between phenotypic profiles is a strong predictor of functional connection between genes and can be used to characterise unannotated genes with known biological pathways (Nichols et al., 2011). Similarly, condition-condition correlations help identify clusters of chemical or environmental perturbations that produce comparable phenotypic effects, enabling researchers to group drugs with similar mechanisms of action or identify synergistic stress responses (Brochado et al., 2018).

The application supports the visualisation of gene-gene and condition-condition correlations (Fig. 6). By using Pearson’s correlation coefficient, ChemGenXplore allows users to assess how gene fitness correlates across various conditions, as well as how conditions correlate based on their impact on phenotype. It provides interactive bar plots that display correlations greater than ± 0.4. Users can adjust the FDR threshold to filter for statistically significant correlations and download the results for further analysis.

**Figure 6.**
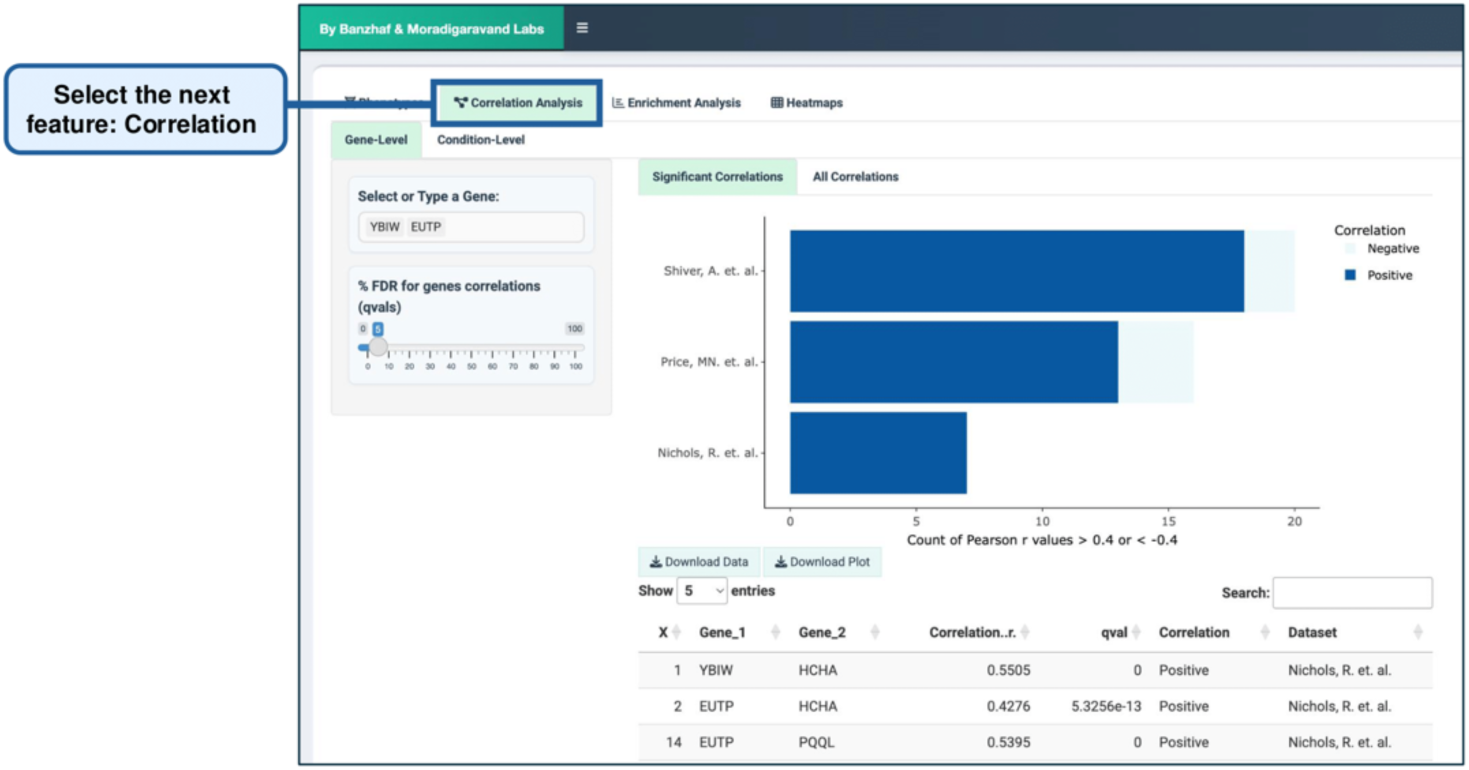
Correlation Analysis. ChemGenXplore supports the visualisation of gene-gene and condition-condition correlations using Pearson’s correlation coefficient. Interactive bar plots display correlations greater than ± 0.4, with an FDR threshold to filter statistically significant correlations. Users can adjust the threshold and download results for further analysis as in the Phenotype Visualisation tab.

### 3.4 Enrichment Analysis

Enrichment analysis identifies biological pathways and functional gene sets associated with phenotypic changes. By integrating GO and KEGG pathway enrichment analysis, ChemGenXplore enables researchers to uncover biological processes, molecular functions, and metabolic pathways enriched in their dataset. Enrichment results are visualised through interactive bar plots and tables, with the ability to filter results by FDR threshold (Fig. 7). This feature helps uncover biological processes and pathways linked to genes of interest. Users can download both the data and plots for further exploration.

**Figure 7.**
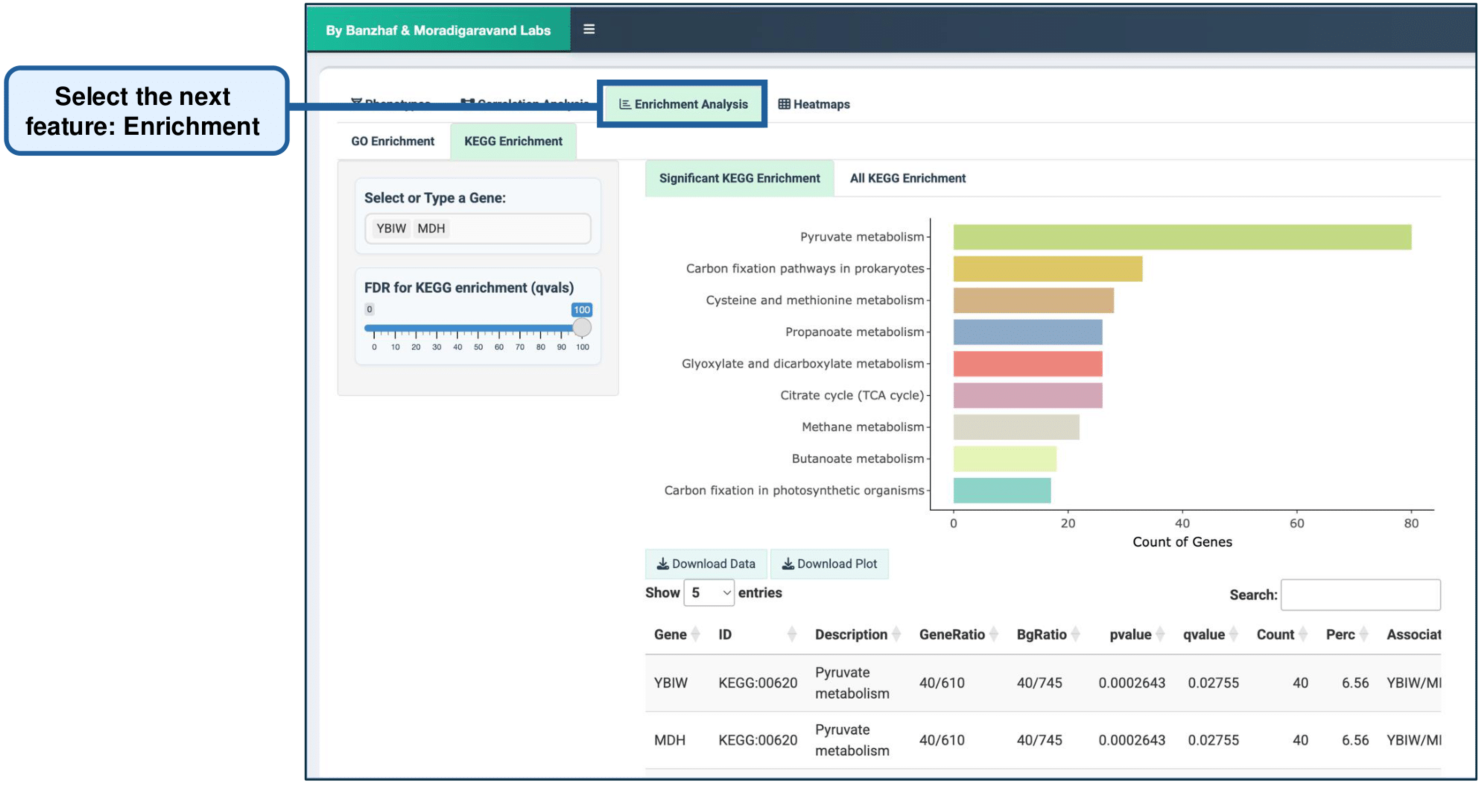
Enrichment Analysis. Users can perform GO and KEGG enrichment analysis, following the same options as in the Phenotype Visualisation and Correlation Analysis tabs.

### 3.5 Interactive Heatmaps

Heatmaps are widely used in chemical genomics, enabling researchers to visualise phenotypic patterns. By clustering genes and conditions based on phenotypic profiles, heatmaps provide an intuitive way to detect functionally related genes. ChemGenXplore allows users to generate customisable, interactive heatmaps to visualise gene-condition fitness data (Fig. 8). Users can select genes and conditions of interest and apply clustering methods to group similar data points. Various clustering algorithms (e.g., complete, single, average, and ward.D) and distance metrics (e.g., Euclidean, Manhattan, etc.) are available, providing flexibility in data exploration. The heatmaps are fully interactive, allowing users to zoom in on specific areas and explore relationships between genes and conditions. Both the heatmap and dendrogram plots are available for download pdf and csv formats.

**Figure 8.**
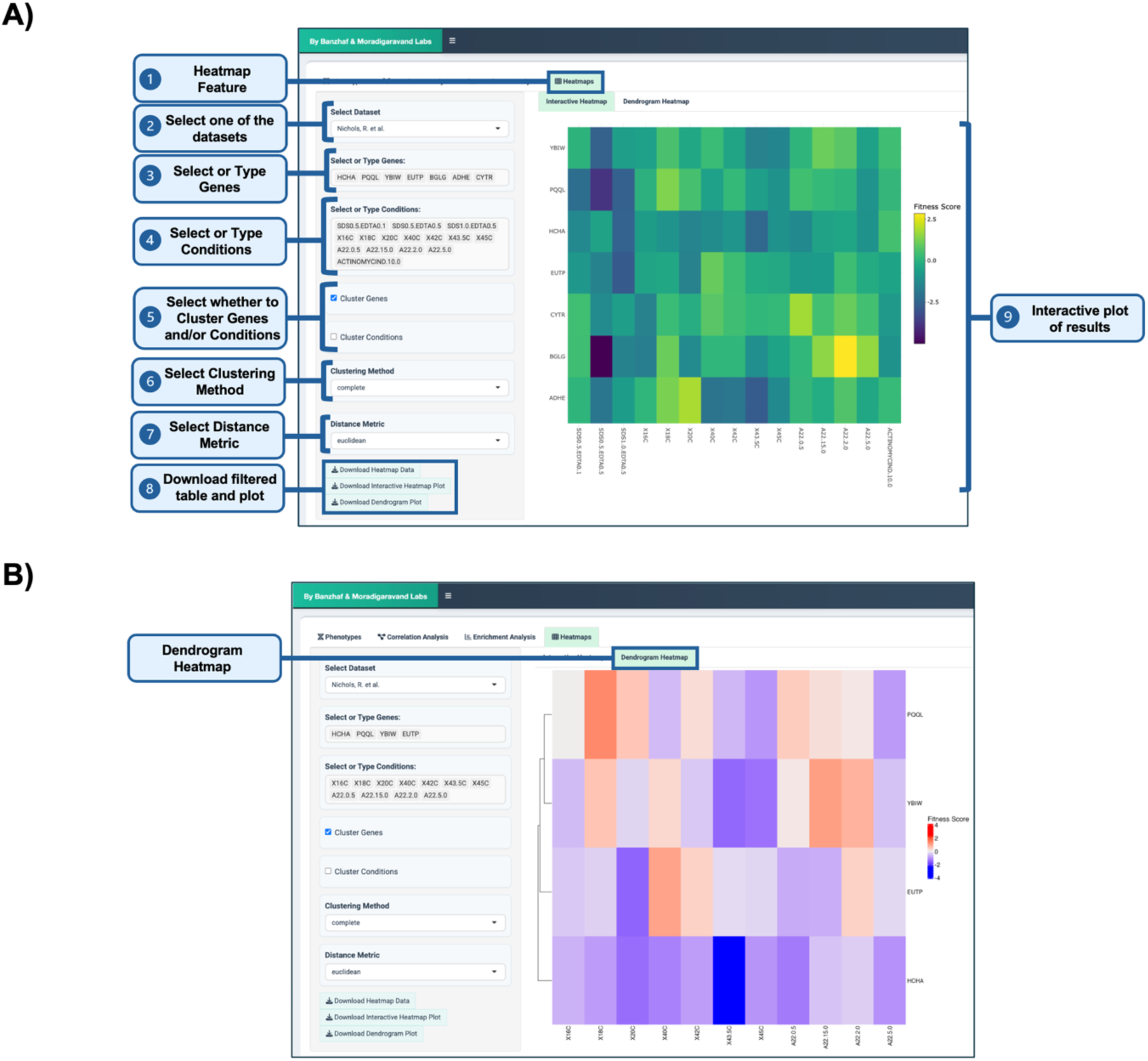
Interactive Heatmaps. **(A)** Users can generate interactive heatmaps to visualise phenotypic relationships and patterns across conditions, applying various clustering methods and distance metrics. **(B)** Corresponding dendrograms display hierarchical relationships between genes and conditions. Both heatmaps and dendrograms are downloadable, following the same options as in the previous tabs.

### 3.6 Case Study: Visualising Antifolate Stress Responses

To demonstrate ChemGenXplore’s utility in uncovering condition-specific phenotypes, we selected a subset of genes involved in tetrahydrofolate (THF) and one-carbon metabolism under antifolate stress. This example replicates a key finding from Nichols et al. (2010), which characterised gene-specific responses to sulfonamides, trimethoprim (TMP), and their combination. The resulting heatmap in Figure 9A reveals that the fitness response of *ΔgcvA*, *ΔgcvP*, *ΔgcvT*, *ΔgcvH*, and *ΔygfA* mutants were selectively reduced under sulfonamides, while Δ*glyA* was sensitive to TMP. *ΔnudB* deletion conferred hypersensitivity to all folate stresses, consistent with its role in folate biosynthesis. In contrast, *ΔfolM* and *ΔfolX* mutants exhibited increased resistance across conditions. These results are consistent with Nichols et al. (2010). Furthermore, we generated a gene–gene correlation network based on Pearson correlations of fitness profiles across antifolate conditions. As shown in Figure 9B, the network reveals clusters of genes with similar responses to antifolate stress. Edge colours indicate the sign of the correlation (positive or negative), while node colours correspond to gene clusters identified using the Louvain algorithm, a modularity-based method for detecting communities within networks (Blondel et al., 2008). This combined visualisation illustrates ChemGenXplore’s ability to uncover similar gene responses and functionally related gene groups under stresses.

**Figure 9.**
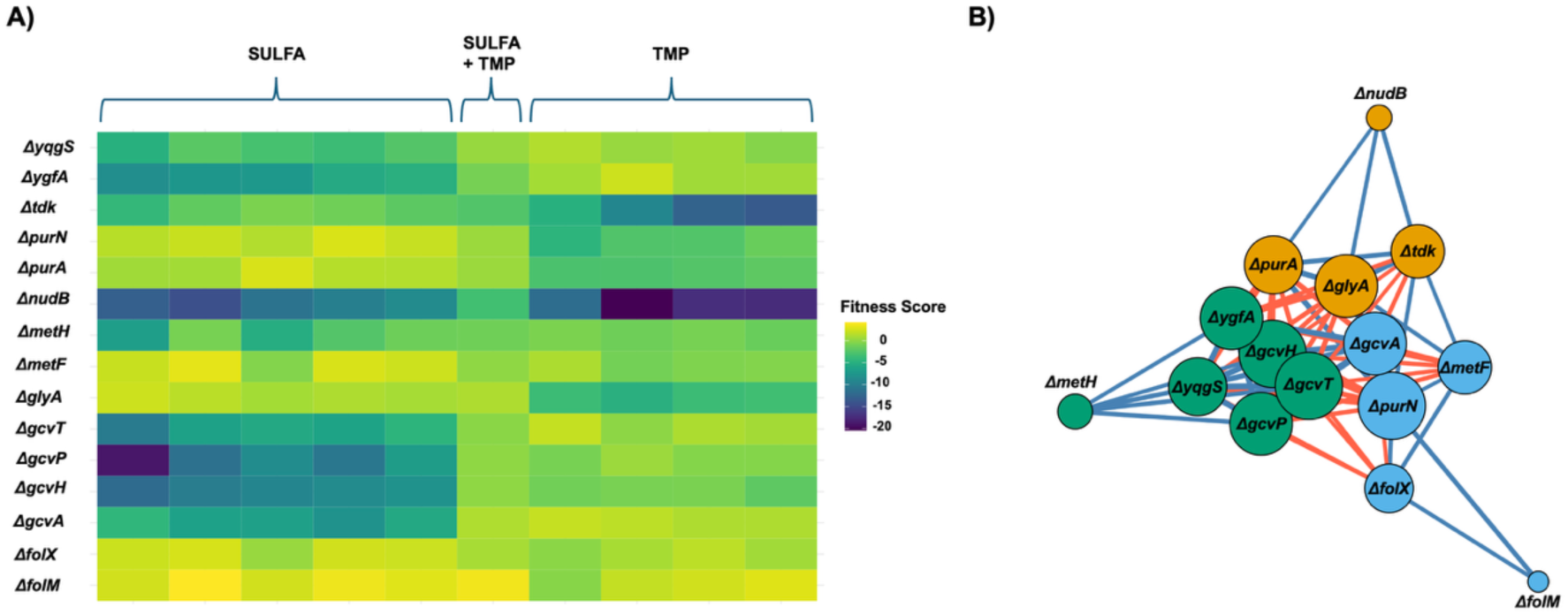
Differential Antifolate Responses Revealed by ChemGenXplore. **A)** A heatmap visualising phenotypic profiles of mutants involved in tetrahydrofolate (THF) and one-carbon metabolism under sulfonamide, trimethoprim (TMP), and combination stress. Clustering reveals distinct responses, including sulfa-specific sensitivity in *Δgcv* and *ΔygfA*, TMP-specific sensitivity in *ΔglyA*, hypersensitivity in *ΔnudB*, and increased resistance in *ΔfolM* and *ΔfolX*. **(B)** Gene–gene correlation network based on Pearson correlation of fitness profiles across antifolate conditions (|r| > 0.6). Blue edges represent positive correlations, red edges indicate negative correlations. Node colours correspond to Louvain clusters.

## 4 Discussion

The first release of ChemGenXplore provides a pre-implemented *Escherichia coli* dataset alongside the option for users to upload their own datasets. In future versions, we aim to expand its functionality by integrating additional species from ongoing and future chemical genomic screens. By consolidating chemical genomic data from various bacterial species, ChemGenXplore serves as a centralised tool for the chemical genomics community, facilitating comparative analyses to identify conserved and species-specific phenotypic responses. A key advantage of this approach is its ability to address a wide range of research questions spanning multiple fields, including gene-function mapping, genetic and drug interactions, and mode-of-action studies for drug discovery. By supporting data sharing, reproducibility, and collaboration, ChemGenXplore will continue to integrate additional datasets and enhance its functionality. We look forward to a growing community of users and contributors to support ongoing research in the field.

## Acknowledgements

We thank members of the Banzhaf and Moynihan labs for thoughtful discussions and feedback. We also gratefully acknowledge King Abdullah University of Science and Technology (KAUST) for providing the web server used to host ChemGenXplore.

## Data availability

The data underlying this article are available in *Zenodo, at <u>DOI: 10.5281/zenodo.15278229</u>.* The datasets were derived from sources in the public domain:

NICHOLS, R. J., SEN, S., CHOO, Y. J., BELTRAO, P., ZIETEK, M., CHABA, R., LEE, S., KAZMIERCZAK, K. M., LEE, K. J., WONG, A., SHALES, M., LOVETT, S., WINKLER, M. E., KROGAN, N. J., TYPAS, A. & GROSS, C. A. 2011. Phenotypic Landscape of a Bacterial Cell. *Cell,* 144, 143-156.

PRICE, M. N., WETMORE, K. M., WATERS, R. J., CALLAGHAN, M., RAY, J., LIU, H., KUEHL, J. V., MELNYK, R. A., LAMSON, J. S., SUH, Y., CARLSON, H. K., ESQUIVEL, Z., SADEESHKUMAR, H., CHAKRABORTY, R., ZANE, G. M., RUBIN, B. E., WALL, J. D., VISEL, A., BRISTOW, J., BLOW, M. J., ARKIN, A. P. & DEUTSCHBAUER, A. M. 2018. Mutant phenotypes for thousands of bacterial genes of unknown function. *Nature,* 557, 503-509.

SHIVER, A. L., OSADNIK, H., KRITIKOS, G., LI, B., KROGAN, N., TYPAS, A. & GROSS, C. A. 2017. Correction: A Chemical-Genomic Screen of Neglected Antibiotics Reveals Illicit Transport of Kasugamycin and Blasticidin S. *PLOS Genetics,* 13, e1006902.

## Funding

This work was supported by the Darwin Trust of Edinburgh. D.M. was supported by the KAUST baseline fund [BAS/1/1108-01-01]. M.B. was supported by a UKRI Future Leaders Fellowship [MR/V027204/1]. DM and GZ were also supported by FCC/1/5932-01-03 from the KAUST Center of Excellence for Smart Health.

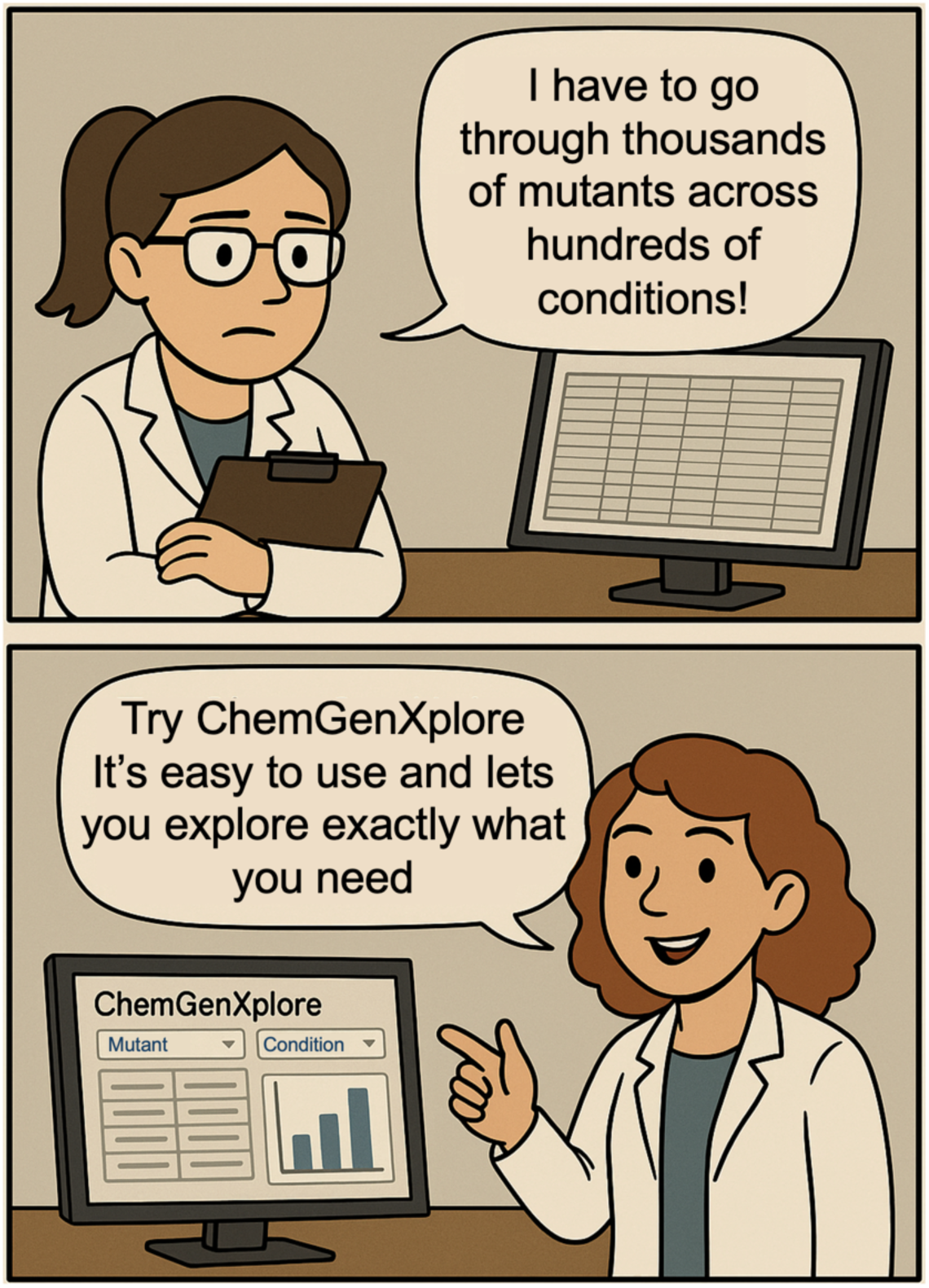

OpenAI. (2025). *Comic generated by ChatGPT* [AI-generated comic]. ChatGPT. https://chat.openai.com

## References

Alodaini, D., Hernandez-Rocamora, V., Boelter, G., Ma, X., ALAO Micheal, B., Doherty Hannah, M., Bryant Jack, A., Moynihan, P., Moradigaravand, D., Glinkowska, M., Vollmer, W. & Banzhaf, M. 2024. Reduced peptidoglycan synthesis capacity impairs growth of E. coli at high salt concentration. mBio, 15, e00325–24.

Arend, D., Lange, M., Pape, J.-M., Weigelt-Fischer, K., Arana-Ceballos, F., Mücke, I., Klukas, C., Altmann, T., Scholz, U. & Junker, A. 2016. Quantitative monitoring of Arabidopsis thaliana growth and development using high-throughput plant phenotyping. Scientific Data, 3, 160055.

Blondel, V. D., Guillaume, J.-L., Lambiotte, R. & Lefebvre, E. 2008. Fast unfolding of communities in large networks. Journal of Statistical Mechanics: Theory and Experiment, 2008, P10008.

Bobonis, J., Mitosch, K., Mateus, A., Karcher, N., Kritikos, G., Selkrig, J., Zietek, M., Monzon, V., Pfalz, B., Garcia-Santamarina, S., Galardini, M., Sueki, A., Kobayashi, C., Stein, F., Bateman, A., Zeller, G., Savitski, M. M., Elfenbein, J. R., Andrews-Polymenis, H. L. & Typas, A. 2022. Bacterial retrons encode phage-defending tripartite toxin–antitoxin systems. Nature, 609, 144–150.

Brochado, A. R., Telzerow, A., Bobonis, J., Banzhaf, M., Mateus, A., Selkrig, J., Huth, E., Bassler, S., Zamarreño Beas, J., Zietek, M., Ng, N., Foerster, S., Ezraty, B., Py, B., Barras, F., Savitski, M. M., Bork, P., Göttig, S. & Typas, A. 2018. Species-specific activity of antibacterial drug combinations. Nature, 559, 259–263.

Brown, J. B., Boley, N., Eisman, R., May, G. E., Stoiber, M. H., Duff, M. O., Booth, B. W., Wen, J., Park, S., Suzuki, A. M., Wan, K. H., Yu, C., Zhang, D., Carlson, J. W., Cherbas, L., Eads, B. D., Miller, D., Mockaitis, K., Roberts, J., Davis, C. A., Frise, E., Hammonds, A. S., Olson, S., Shenker, S., Sturgill, D., Samsonova, A. A., Weiszmann, R., Robinson, G., Hernandez, J., Andrews, J., Bickel, P. J., Carninci, P., Cherbas, P., Gingeras, T. R., Hoskins, R. A., Kaufman, T. C., Lai, E. C., Oliver, B., Perrimon, N., Graveley, B. R. & Celniker, S. E. 2014. Diversity and dynamics of the Drosophila transcriptome. Nature, 512, 393–9.

Cain, A. K., Barquist, L., Goodman, A. L., Paulsen, I. T., Parkhill, J. & VAN Opijnen, T. 2020. A decade of advances in transposon-insertion sequencing. Nature Reviews Genetics, 21, 526–540.

Chen, D., Fu, L.-Y., Hu, D., Klukas, C., Chen, M. & Kaufmann, K. 2018. The HTPmod Shiny application enables modeling and visualization of large-scale biological data. Communications Biology, 1, 89.

Collins, S. R., Schuldiner, M., Krogan, N. J. & Weissman, J. S. 2006. A strategy for extracting and analyzing large-scale quantitative epistatic interaction data. Genome Biology, 7, R63.

De Hoon, M. J. L., Imoto, S., Nolan, J. & Miyano, S. 2004. Open source clustering software. Bioinformatics, 20, 1453–1454.

Dixon, S. J., Costanzo, M., Baryshnikova, A., Andrews, B. & Boone, C. 2009. Systematic mapping of genetic interaction networks. Annu Rev Genet, 43, 601–25.

Doherty, H. M., Kritikos, G., Galardini, M., Banzhaf, M. & Moradigaravand, D. 2023. ChemGAPP: a tool for chemical genomics analysis and phenotypic profiling. Bioinformatics, 39.

Fajardo, A., Martínez-Martín, N., Mercadillo, M., Galán, J. C., Ghysels, B., Matthijs, S., Cornelis, P., Wiehlmann, L., Tümmler, B., Baquero, F. & Martínez, J. L. 2008. The Neglected Intrinsic Resistome of Bacterial Pathogens. PLOS One, 3, e1619.

French, S., Mangat, C., Bharat, A., Cote, J. P., Mori, H. & Brown, E. D. 2016. A robust platform for chemical genomics in bacterial systems. Mol Biol Cell, 27, 1015–25.

Gerstein, M. B., Rozowsky, J., Yan, K.-K., Wang, D., Cheng, C., Brown, J. B., Davis, C. A., Hillier, L., Sisu, C., Li, J. J., Pei, B., Harmanci, A. O., Duff, M. O., Djebali, S., Alexander, R. P., Alver, B. H., Auerbach, R., Bell, K., Bickel, P. J., Boeck, M. E., Boley, N. P., Booth, B. W., Cherbas, L., Cherbas, P., Di, C., Dobin, A., Drenkow, J., Ewing, B., Fang, G., Fastuca, M., Feingold, E. A., Frankish, A., Gao, G., Good, P. J., Guigó, R., Hammonds, A., Harrow, J., Hoskins, R. A., Howald, C., Hu, L., Huang, H., Hubbard, T. J. P., Huynh, C., Jha, S., Kasper, D., Kato, M., Kaufman, T. C., Kitchen, R. R., Ladewig, E., Lagarde, J., Lai, E., Leng, J., Lu, Z., Maccoss, M., May, G., Mcwhirter, R., Merrihew, G., Miller, D. M., Mortazavi, A., Murad, R., Oliver, B., Olson, S., Park, P. J., Pazin, M. J., Perrimon, N., Pervouchine, D., Reinke, V., Reymond, A., Robinson, G., Samsonova, A., Saunders, G. I., Schlesinger, F., Sethi, A., Slack, F. J., Spencer, W. C., Stoiber, M. H., Strasbourger, P., Tanzer, A., Thompson, O. A., Wan, K. H., Wang, G., Wang, H., Watkins, K. L., Wen, J., Wen, K., Xue, C., Yang, L., Yip, K., Zaleski, C., Zhang, Y., Zheng, H., Brenner, S. E., Graveley, B. R., Celniker, S. E., Gingeras, T. R. & Waterston, R. 2014. Comparative analysis of the transcriptome across distant species. Nature, 512, 445–448.

Gomez Maria, J. & Neyfakh Alexander, A. 2006. Genes Involved in Intrinsic Antibiotic Resistance of Acinetobacter baylyi. Antimicrobial Agents and Chemotherapy, 50, 3562–3567.

Hayward, R. J., Ebbecke, T., Fricke, H., Nguyen, V. Q. & Barquist, L. 2025. Micromix: web infrastructure for visualizing and remixing microbial ’omics data. Gigascience, 14.

Houle, D., Govindaraju, D. R. & Omholt, S. 2010. Phenomics: the next challenge. Nature Reviews Genetics, 11, 855–866.

Koo, B.-M., Kritikos, G., Farelli, J. D., Todor, H., Tong, K., Kimsey, H., Wapinski, I., Galardini, M., Cabal, A., Peters, J. M., Hachmann, A.-B., Rudner, D. Z., Allen, K. N., Typas, A. & Gross, C. A. 2017. Construction and Analysis of Two Genome-Scale Deletion Libraries for <EM>Bacillus subtilis</EM>. Cell Systems, 4, 291–305.e7.

Kritikos, G., Banzhaf, M., Herrera-Dominguez, L., Koumoutsi, A., Wartel, M., Zietek, M. & Typas, A. 2017. A tool named Iris for versatile high-throughput phenotyping in microorganisms. Nature Microbiology, 2, 17014.

Nichols, R. J., Sen, S., Choo, Y. J., Beltrao, P., Zietek, M., Chaba, R., Lee, S., Kazmierczak, K. M., Lee, K. J., Wong, A., Shales, M., Lovett, S., Winkler, M. E., Krogan, N. J., Typas, A. & Gross, C. A. 2011. Phenotypic Landscape of a Bacterial Cell. Cell, 144, 143–156.

Price, M. N., Wetmore, K. M., Waters, R. J., Callaghan, M., Ray, J., Liu, H., Kuehl, J. V., Melnyk, R. A., Lamson, J. S., Suh, Y., Carlson, H. K., Esquivel, Z., Sadeeshkumar, H., Chakraborty, R., Zane, G. M., Rubin, B. E., Wall, J. D., Visel, A., Bristow, J., Blow, M. J., Arkin, A. P. & Deutschbauer, A. M. 2018. Mutant phenotypes for thousands of bacterial genes of unknown function. Nature, 557, 503–509.

Saldanha, A. J. 2004. Java Treeview—extensible visualization of microarray data. Bioinformatics, 20, 3246–3248.

Schadt, E. E., Turner, S. & Kasarskis, A. 2010. A window into third-generation sequencing. Human Molecular Genetics, 19, R227–R240.

Schuldiner, M., Collins, S. R., Thompson, N. J., Denic, V., Bhamidipati, A., Punna, T., Ihmels, J., Andrews, B., Boone, C., Greenblatt, J. F., Weissman, J. S. & Krogan, N. J. 2005. Exploration of the Function and Organization of the Yeast Early Secretory Pathway through an Epistatic Miniarray Profile. Cell, 123, 507–519.

Shiver, A. L., Osadnik, H., Kritikos, G., Li, B., Krogan, N., Typas, A. & Gross, C. A. 2017. Correction: A Chemical-Genomic Screen of Neglected Antibiotics Reveals Illicit Transport of Kasugamycin and Blasticidin S. PLOS Genetics, 13, e1006902.

Sullivan, Alessandra M., Arsovski, Andrej A., Lempe, J., Bubb, Kerry L., Weirauch, Matthew T., Sabo, Peter J., Sandstrom, R., Thurman, Robert E., Neph, S., Reynolds, Alex P., Stergachis, Andrew B., Vernot, B., Johnson, Audra K., Haugen, E., Sullivan, Shawn T., Thompson, A., Neri, Fidencio V., Iii, Weaver, M., Diegel, M., Mnaimneh, S., Yang, A., Hughes, Timothy R., Nemhauser, Jennifer L., Queitsch, C. & Stamatoyannopoulos, JOHN A. 2014. Mapping and Dynamics of Regulatory DNA and Transcription Factor Networks in <EM>A.&#xa0;thaliana</EM>. Cell Reports, 8, 2015–2030.

Tamae, C., Liu, A., Kim, K., Sitz, D., Hong, J., Becket, E., Bui, A., Solaimani, P., Tran Katherine, P., Yang, H. & Miller Jeffrey, H. 2008. Determination of Antibiotic Hypersensitivity among 4,000 Single-Gene-Knockout Mutants of Escherichia coli. Journal of Bacteriology, 190, 5981–5988.

Tong, A. H., Evangelista, M., Parsons, A. B., Xu, H., Bader, G. D., Pagé, N., Robinson, M., Raghibizadeh, S., Hogue, C. W., Bussey, H., Andrews, B., Tyers, M. & Boone, C. 2001. Systematic genetic analysis with ordered arrays of yeast deletion mutants. *Science (New York*, N.Y.), 294, 2364–2368.

Tsankov, A. M., Gu, H., Akopian, V., Ziller, M. J., Donaghey, J., Amit, I., Gnirke, A. & Meissner, A. 2015. Transcription factor binding dynamics during human ES cell diterentiation. Nature, 518, 344–9.

Typas, A., Banzhaf, M., Van Den Berg Van Saparoea, B., Verheul, J., Biboy, J., Nichols, R. J., Zietek, M., Beilharz, K., Kannenberg, K., VON Rechenberg, M., Breukink, E., DEN Blaauwen, T., Gross, C. A. & Vollmer, W. 2010. Regulation of Peptidoglycan Synthesis by Outer-Membrane Proteins. Cell, 143, 1097–1109.

Williams, G., Ahmad, H., Sutherland, S., Haycocks, J., Benedict, S., Hart, A., Doherty, H., Sullivan, R., Alao, M., Ma, X., Xu, Q., Bryant, J., Glinkowska, M., Banks, P., Moynihan, P., Milner, M., Moradigaravand, D. & Banzhaf, M. 2025. High-Throughput Chemical Genomic Screening: A Step-by-Step Workflowfrom Plate to Phenotype. Preprints. Preprints.

